# Drivers of antimicrobial resistance amongst intestinal *Escherichia coli* isolated from children in South Asia and sub-Saharan Africa

**DOI:** 10.1101/233460

**Authors:** Danielle J. Ingle, Myron M. Levine, Karen L. Kotloff, Kathryn E. Holt, Roy M. Robins-Browne

**Author notes:** Current address: National Centre for Epidemiology & Population Health, The Australian National University, and Microbiological Diagnostic Unit Public Health Laboratory, Department of Microbiology and Immunology, Peter Doherty Institute for Infection and Immunity, The University of Melbourne, Victoria, Australia (Danielle J. Ingle). These authors contributed equally to this work (Kathryn E. Holt, Roy M. Robins-Browne). Corresponding author: (RMRB).

## Abstract

Antimicrobial resistance (AMR) dynamics are poorly understood in developing countries, where data on the prevalence of AMR in enteric bacteria are sparse, particularly among children and in the community setting. Here we use a combination of phenotyping, genomics and antimicrobial usage data to investigate patterns of AMR amongst atypical enteropathogenic *E. coli* (aEPEC) strains isolated from children <5 years old in seven countries (four in sub-Saharan Africa and three in South Asia) over a three-year period. We detected very high rates of AMR, with 65% of isolates displaying resistance to ≥3 drug classes; the rates of AMR were the same amongst strains associated with diarrhea and strains that were carried asymptomatically. Whole genome sequencing identified a diversity of genetic mechanisms for AMR, which could explain >95% of observed phenotypic resistance. Analysis of AMR gene co-occurrence revealed clusters of acquired AMR genes that were frequently co-located on small plasmids and transposons, providing opportunities for acquisition of multidrug resistance in a single step. We used discriminant analysis to investigate potential drivers of AMR within the bacterial population, and found that genetic determinants of AMR were associated with geographical location of isolation but not with phylogenetic lineage of the *E. coli* strain or disease status of the human host. Comparison with antimicrobial usage data showed that the prevalence of resistance to newer drugs (fluoroquinolones and third-generation cephalosporins) was correlated with usage, which was generally higher in South Asia than Africa. In particular, fluoroquinolone resistance-associated mutations in *gyrA* were significantly associated with use of these drugs for treatment of diarrheic children. Notably resistance to older drugs such as trimethoprim, chloramphenicol and ampicillin, which are conferred by acquired AMR genes that were frequently clustered together in mobile genetic elements, were common in all locations despite differences in usage; this suggests that reversion to sensitivity is unlikely to occur even if these drugs are removed from circulation. This study provides much-needed insights into the frequencies of AMR in intestinal *E. coli* in community-based children in developing countries and to antimicrobial usage for diarrhea where the burden of infections is greatest.

## Introduction

Certain pathotypes of *Escherichia coli* (*E. coli*) are a leading cause of diarrhea in children, especially in the developing countries of sub-Saharan Africa and South Asia^1^. Pathogenic varieties of *E. coli* have evolved on multiple occasions; atypical enteropathogenic *E. coli* (aEPEC) is one such pathotype, which is defined by the presence of the locus of enterocyte effacement (LEE) pathogenicity island, and the absence of Shiga toxins (denoting enterohemorrhagic *E. coli*) and type IV bundle-forming pili (denoting typical EPEC)^2-5^. We have recently identified distinct lineages of aEPEC and ten common clonal groups (CGs)^6^. Importantly, unlike most other *E. coli* pathotypes, aEPEC has various animal reservoirs and, while it can cause a range of disease symptoms in humans from sporadic to persistent diarrhea, it may also colonize the human gut asymptomatically^1,7-9^.

Antimicrobial resistance (AMR) has been reported in *E. coli* from multiple animal sources, the environment as well as in hospital-acquired infections globally^10-15^. Of greatest concern are multi-drug resistant (MDR) *E. coli* (non-susceptible to one or more agents in at least three antimicrobial categories)^16^, particularly those that are fluoroquinolone resistant and/or produce extended-spectrum beta-lactamases (ESBL). While several recent studies have reported increases in ESBLs ^17,18^, gentamicin ^19,20^, ciprofloxacin^20-24^ and revealed diverse AMR profiles, these data have largely come from *E. coli* responsible for disease in hospital settings. Thus, there remain significant gaps in knowledge of the prevalence of AMR in human intestinal *E. coli* populations globally, particularly in the developing nations of sub-Saharan Africa and South Asia^25^.

The rise of AMR, and particularly MDR in bacteria, is a significant public health concern as it makes infections non-responsive to treatment while increasing the reservoir of resistance-encoding genes. Although antimicrobials are not recommended for the treatment of uncomplicated gastroenteritis, they are commonly administered to diarrheic children in developing countries to prevent or treat dysentery^26^ and prolonged diarrhea, of which aEPEC is a major cause^8,9^. Enhancing our knowledge of AMR amongst gut-dwelling *E. coli* has broader implications: first, because *E. coli* is a leading cause of extra-intestinal infections in hospitals, and strains colonizing the gastrointestinal tract of patients are the major reservoir of these infections; and second, because most AMR in *E. coli* is encoded on mobile genetic elements that are subject to HGT, which enable the rapid dissemination of genes that prevent susceptibility to drugs and will be maintained under positive selective pressure^27,28^. Hence AMR determinants may be shared between *E. coli* strains regardless of pathotype. For these reasons, it is useful to determine the AMR profiles of intestinal *E. coli*, such as aEPEC, as these can provide an indication of the prevalence of AMR in Enterobacteriaceae in general and the gut microbiome as a whole.

Here we present AMR data for 185 aEPEC isolates collected as part of the Global Enteric Multicenter Study (GEMS)^1,29^, using phenotypic susceptibility data and whole genome sequence analysis to identify the mechanisms of resistance and potential drivers of variation in AMR profiles. These isolates, collected from healthy children in the community and children with diarrhea at seven sites in sub-Saharan Africa and South Asia, provide a unique opportunity to investigate the prevalence of AMR in a bacterial population that was not selected on the basis of AMR profile.

## Results

### Antimicrobial susceptibility profiles

Our collection of 185 aEPEC isolates (Table S1) was tested for susceptibility to 16 antimicrobials (Table S2). Resistance was detected to 14 of the drugs investigated (Fig 1A; Table S3), and 121 isolates (65%) were MDR (i.e., resistant to ≥3 drug classes) (Fig 1B). No resistance was detected to amikacin (an aminoglycoside) or meropenem (a carbapenem). Only 35 isolates (19%) were susceptible to all drugs tested (17 from cases and 18 from controls). Resistance to ‘older’ antimicrobials was common, with 121 (65%) of isolates resistant to ampicillin, 80 (43%) resistant to streptomycin, 124 (67%) resistant to trimethoprim, 122 (66%) resistant to trimethoprim/sulfamethoxazole and 104 (56%) resistant to tetracycline (Fig 1A). Around half (n=96, 52%) of isolates were resistant to three or more of these first-line drugs. While streptomycin resistance was common, 180 isolates (97%) were susceptible to all other aminoglycosides tested; the exceptions being one isolate from India that was resistant to tobramycin and intermediately susceptible to gentamicin; and four isolates from India that were resistant to both drugs. Fluoroquinolone resistance was relatively infrequent, with 31 (17%) isolates resistant to norfloxacin and 8 (4%) resistant to both norfloxacin and ciprofloxacin. Resistance to chloramphenicol (11%) and azithromycin (7%) was also infrequent. Amongst the beta-lactam antibiotics, while ampicillin resistance was common (65%), resistance to the cephalosporins, ceftriaxone (third generation; 3%), ceftazidime (third generation; 3%) and cefepime (fourth generation; 1.6%) was rare (Fig 1A-B).

**Fig 1.**
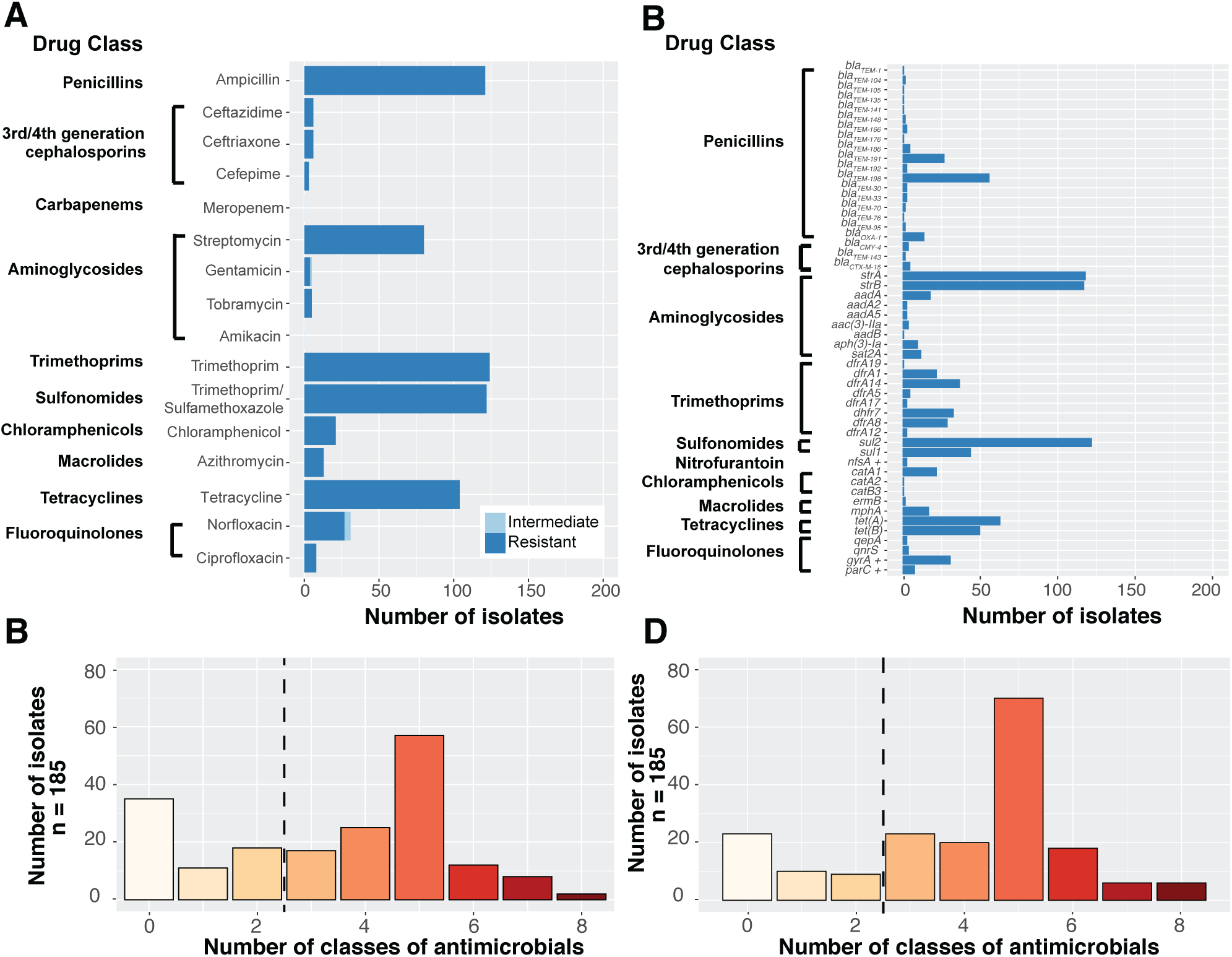
Prevalence of AMR-associated gene content and AMR phenotypes in 185 aEPEC isolates. Panels (A) and (B) summarize the AMR phenotypes of the aEPEC isolates. (A) AMR profiles are grouped by drug class. (B) Histogram of number of drug classes to which aEPEC strains were phenotypically resistant. Panels (C) and (D) summarize AMR-associated gene content in aEPEC strains. (C) Genes detected in the genomes associated with antimicrobial resistance are shown to the left of the graphs and are grouped by drug class. + indicates genes which have point mutations resulting in antimicrobial resistance and are not acquired through horizontal gene transfer. (D) Histogram of number of classes to which aEPEC strains were detected as having AMR-associated gene content.

### Genetic determinants of AMR

The genomes of the 185 aEPEC isolates were screened for known genetic determinants of AMR, including acquired genes as well as point mutations in chromosomal genes associated with resistance to fluoroquinolones and nitrofurantoin (Fig 1C; Fig S1 and Table S4). Thirty acquired AMR genes were detected, along with four point mutations (two in *gyrA*, one in *parC*, and one in *nfsA).* Four acquired AMR genes were detected in more than half the isolates: *bla*_TEM_ genes (ampicillin), *strA* and *strB* (streptomycin), and *sul2* (sulfonamides). Alleles of dihydrofolate reductase (*dfr*) genes encoding trimethoprim resistance were detected in 132 (71%) isolates. The most common of these were *dfrA1* (12%), *dfrA5* (23%), *dfrA7* (20%), and *dfrA8* (16%). The common AMR genes were determined to be predominately co-mobilized on a small number of common mobile elements (Fig S2; Fig S3 and Supplementary Results). Extensive diversity of AMR genotypes was observed, with 104 distinct combinations of AMR determinants across the 185 isolates (Fig S1).

### Prediction of AMR phenotypes from genotypes

Phenotypic resistance profiles were largely explained by known genetic determinants of AMR (Fig 1, Table 1). For most drugs, the detection of known genetic determinants was both sensitive (>95%) and specific (>90%) for AMR phenotypes. The frequency of very major errors (failing to detect resistance when it is present) exceeded the minimum acceptable threshold of 1.5% for five drugs (Table 1). These were: ampicillin (4.9%); streptomycin (2.2%); trimethoprim (2.2%); trimethoprim/sulfamethoxazole (2.2%) and tetracycline (2.2%). Major errors (predicting resistance when none is present) were also detected for these and several other antimicrobials (Table 1). The highest major error rates were observed for streptomycin (26.5%), ampicillin (7.0%), trimethoprim (5.4%), and tetracycline (4.3%). The sensitivity, specificity, positive predictive value (PPV) and negative predictive value (NPV) for each drug are summarized in Table S5. Sensitivity and NPV were greater than 90% for all drugs tested with the exception of azithromycin (85% sensitivity) and ampicillin 853% NPV). Specificity and PPV were more variable, reflecting the error rates outlined above (Table 1).

**Table 1.** Comparison of phenotypic and genotypic AMR profiles of 185 aEPEC

|  | Resistant phenotype | Resistant genotype | False susceptible<sup>^</sup> | Susceptible phenotype | Susceptible genotype | False resistant<sup>^^</sup> |
| --- | --- | --- | --- | --- | --- | --- |
| **Beta-lactams** |  |  |  |  |  |  |
| Ampicillin | 121 | 112 | **9 (4.9%)** | 64 | 51 | **13 (7.0%)** |
| Cefepime | 3 | 3 | 0 | 182 | 178 | 4 (2.2%) |
| Ceftriaxone | 6 | 6 | 0 | 179 | 174 | 5 (2.7%) |
| Ceftazidime | 6 | 6 | 0 | 179 | 174 | 5 (2.7%) |
| Meropenem | 0 | 0 | 0 | 185 | 185 | 0 |
| **Aminoglycosides** |  |  |  |  |  |  |
| Streptomycin | 80 | 76 | **4 (2.2%)** | 105 | 56 | **49 (26.5%)** |
| Gentamicin | 5 | 5 | 0 | 180 | 180 | 0 |
| Tobramycin | 5 | 5 | 0 | 180 | 180 | 0 |
| Amikacin | 0 | 0 | 0 | 185 | 185 | 0 |
| **Folate pathway inhibitors** |  |  |  |  |  |  |
| Trimethoprim | 124 | 120 | **4 (2.2%)** | 61 | 50 | 11 (0.9%) |
| Trimethoprim/<br>sulfamethoxazole* | 122 | 118 | **4 (2.2%)** | 63 | 53 | **10 (5.4%)** |
| **Nitrofurantoin** |  |  |  |  |  |  |
| Nitrofurantoin | 0 | 0 | 0 | 185 | 182 | 3 (1.6%) |
| **Chloramphenicol** |  |  |  |  |  |  |
| Chloramphenicol | 21 | 19 | 2 (1.1%) | 164 | 160 | 4 (2.2%) |
| **Macrolides** |  |  |  |  |  |  |
| Azithromycin | 13 | 11 | 2 (1.1%) | 172 | 166 | **6 (3.2%)** |
| **Tetracyclines** |  |  |  |  |  |  |
| Tetracycline | 104 | 100 | **4 (2.2%)** | 81 | 73 | **8 (4.3%)** |
| **Fluoroquinolones** |  |  |  |  |  |  |
| Ciprofloxacin** | 8 | 8 | 0 | 177 | 173 | 4 (2.2%) |
| Norfloxacin | 31 | 30 | 1 (0.5%) | 154 | 149 | 5 (2.7%) |

\* When two genes are required for resistance, both were required for genotypic resistance: *strA* and *strB* for streptomycin, and a *dfr* gene plus a *sul* gene for resistance to trimethoprim/sulfamethoxazole
\*\*At least one point mutation in *gyrA* and a second in *gyrA*, *gyrB*, *parC* or *parE*
<sup>^</sup> very major errors of >1.5% shown in boldface font
<sup>^^</sup> major errors of >3.0% shown in boldface font

### Sources of variation in AMR profiles

Given the diversity of AMR profiles observed in our aEPEC collection, we sought to determine if the distribution of AMR genes was associated with disease status, phylogenetic lineage, or geographic location where the strain was isolated. First we compared the frequency of AMR phenotypes and genotypes amongst aEPEC isolated from diarrhea cases and asymptomatic controls. Only data from confirmed cases (n=94) and controls (n=88) were used for this analysis. For each drug class, neither AMR phenotype, nor AMR predicted from genotype, was statistically different between cases and controls (Table S6). Also, the frequencies of individual AMR genes in isolates from case and controls were similar (Fig 2A). As AMR determinants were equally distributed amongst cases and controls, all isolates were pooled together for further analysis of lineage and region.

**Fig 2.**
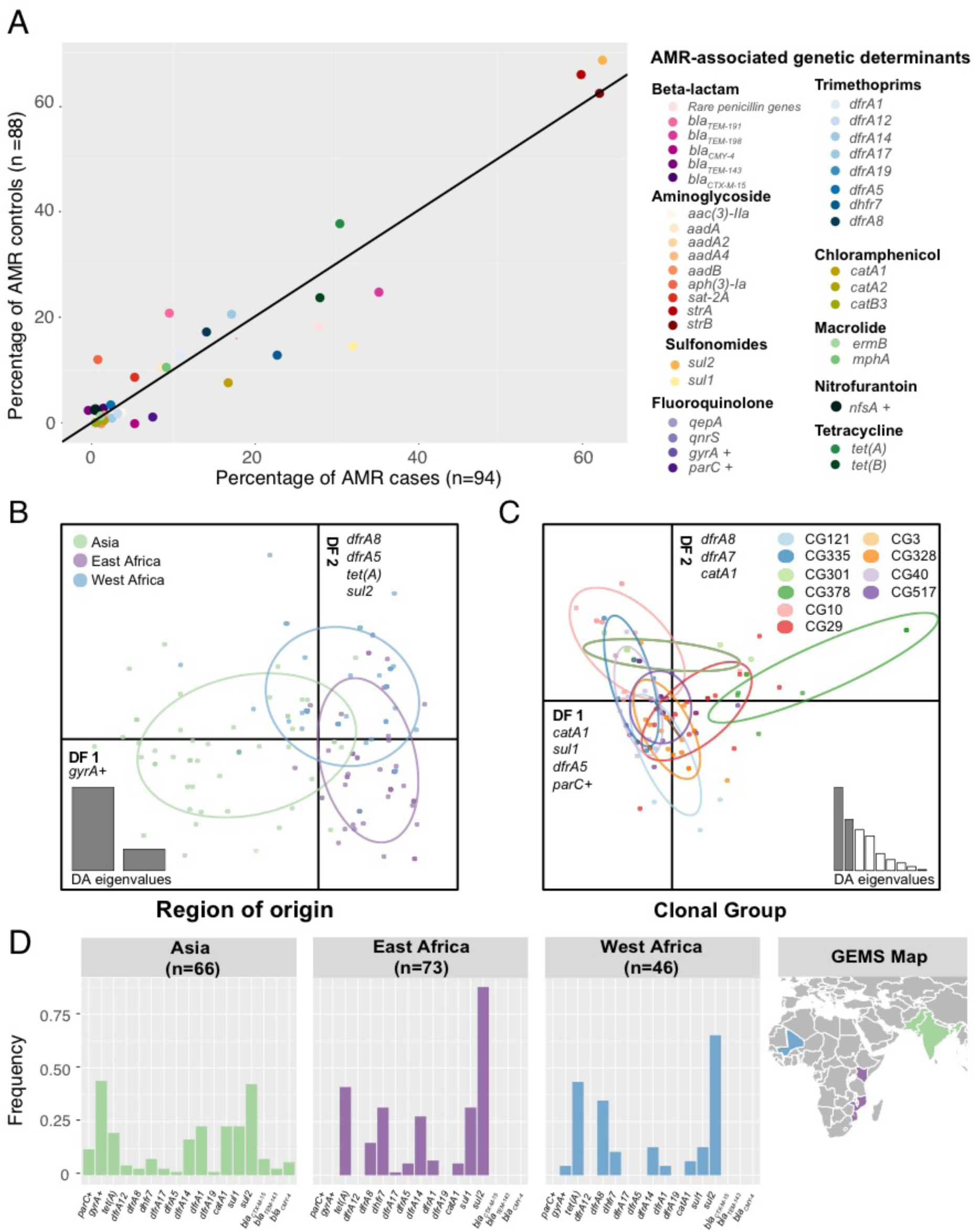
AMR gene content is explained by region of isolation not disease status or clonal group. (A) Frequencies of AMR-associated genes in aEPEC by case or control status. Genes encoding AMR are shown to the right of the graph and are grouped by drug class. (B–C) Discriminant analysis of principal components (DAPC) based on known genetic determinants of AMR. The graphs display the discriminant functions (DF) that best discriminate isolates into region of isolation (B) or clonal group (C). Data points are colored based on their demographic group (according to inset legend) and the genetic determinants most correlated with the DFs are labeled on the DF axes. (D) Frequency of AMR genetic determinants that differed between Asia, East Africa and West Africa. + indicates these genes have point mutations resulting in AMR and are not acquired through horizontal gene transfer.

Discriminant analysis of principal components (DAPC)^30^ on the binary matrix of AMR genetic determinants (Table S4; Fig 3B-C) revealed that the first 15 PCs accounted for >93% of the variance in AMR profiles, and were retained for discriminant analysis by phylogenetic lineage (clonal groups, defined in Ingle *et al.* ^6^ and highlighted in Fig S1) or by region of origin (East Africa, West Africa and Asia; see Fig 2 and Fig S4)).

**Fig 3.**
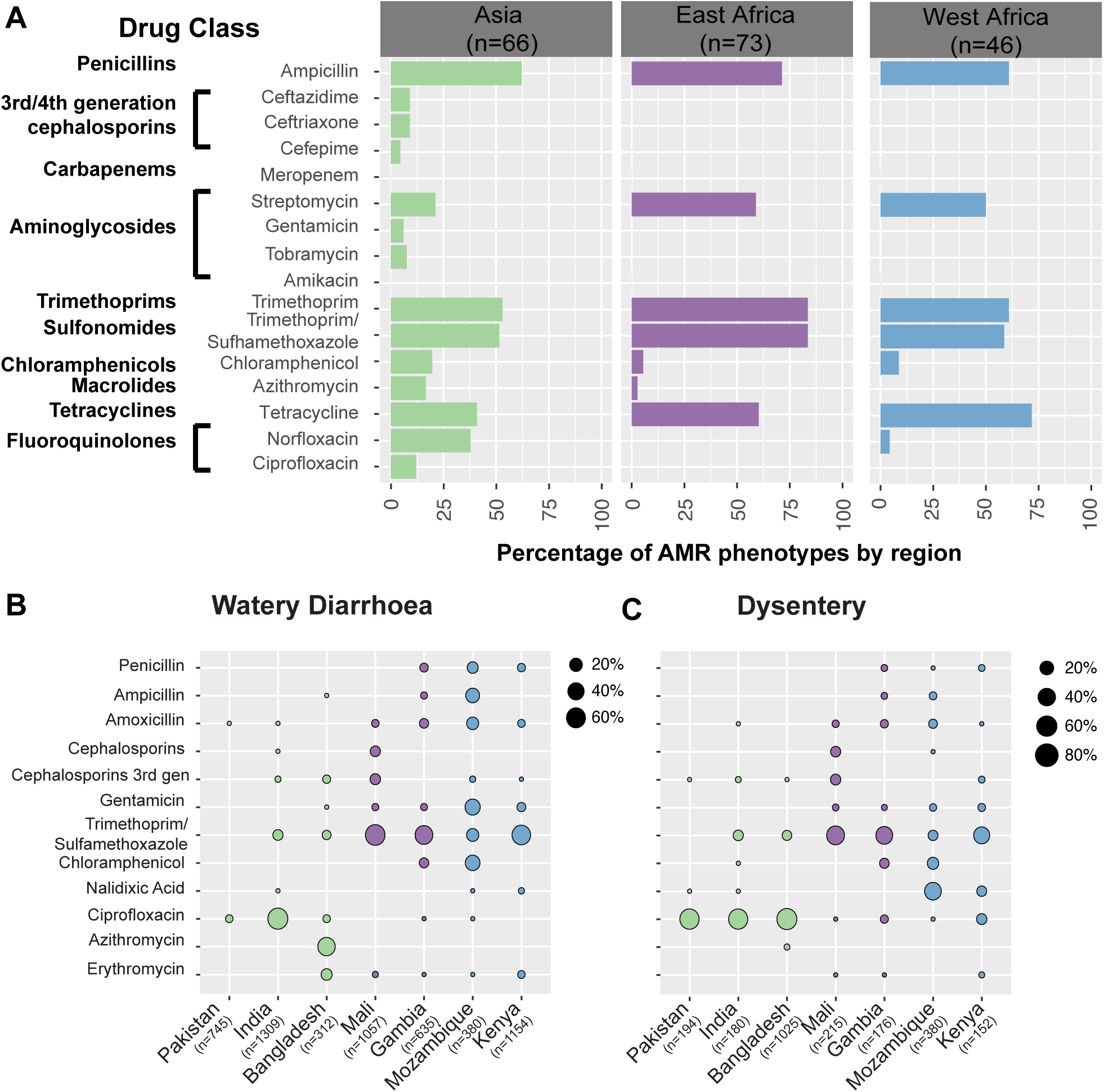
AMR phenotypes by region and antimicrobial use at study sites. (B) AMR phenotypes of GEMS isolates stratified by region of isolation. (B-C) Percentage of antimicrobials prescribed to patients with watery diarrhea (B) or dysentery (C) at each of the seven sites.

Variation in AMR determinants were not associated with clonal group, except for CG378 which was characterized by a lack of most of the common AMR genes (*bla*_TEM_ variants, *sul2*, *tet(A)* and *tetB*) and the presence of the rarer *catA* gene (present in 7 of 9 CG378 isolates compared to 12.5% of non-CG378 isolates; OR=24, *P* <10^−4^). Variation in AMR determinants was associated with region of origin (Fig 2B, Fig S4B). Discriminant function 1 (DF1) separated Asian from African isolates and was associated with *gyrA* SNPs; DF2 separated East from West African isolates and was associated with *dfrA8*, *dfrA5*, *tet(A)* and *sul2*. The various *dfrA* genes were differentially distributed across the various GEMS sites (Fig 2D). For example, *dfrA1* predominated at Asian sites and *dfrA8* was commonest at West African sites whilst *dfrA14* and *dhfr7* were common in Mozambique and Kenya, respectively (Fig 2D). Further, *tet(A)* was more common at West and East African sites than in Asia. These genetic differences were reflected in AMR phenotypes, as resistance to ciprofloxacin and third-generation cephalosporins was identified only in strains from Asia, while resistance to tetracycline was more common in African than in Asian isolates (Fig 3A).

### Differences in local antimicrobial drug usage

We hypothesized that regional differences in AMR and associated genetic determinants may reflect differences in local selective pressure from antimicrobial exposure. The broad patterns of antimicrobial use for the treatment of diarrhea across the study sites showed that trimethoprim/sulfamethoxazole and penicillins were used more frequently at African sites, while macrolides (azithromycin) and fluoroquinolones (in particular ciprofloxacin) were used more frequently in Asia (Fig 3B-C). These patterns of antimicrobial usage showed some association with AMR phenotypes insofar as we observed higher levels of both usage and resistance for azithromycin and fluoroquinolones at Asian sites, and for trimethoprim at East African sites (Fig 3). We could not formally test these associations, however, due to the small numbers of observations at most regional subgroups and the limited variation in usage of most drugs across sites. We therefore investigated the associations between usage and resistance for the two drugs that showed substantial (>10%) usage at three or more study sites; these were ciprofloxacin and trimethoprim. Across the seven sites, ciprofloxacin usage was significantly associated with the prevalence of substitutions in the QRDR of *gyrA* and*parC* (R^2^ = 0.87, *P* = 0.002; Fig 4). By contrast, trimethoprim usage was not associated with the prevalence of horizontally acquired *dfr* genes that confer resistance to the drug (R^2^=0.04, P >0.5).

**Fig 4.**
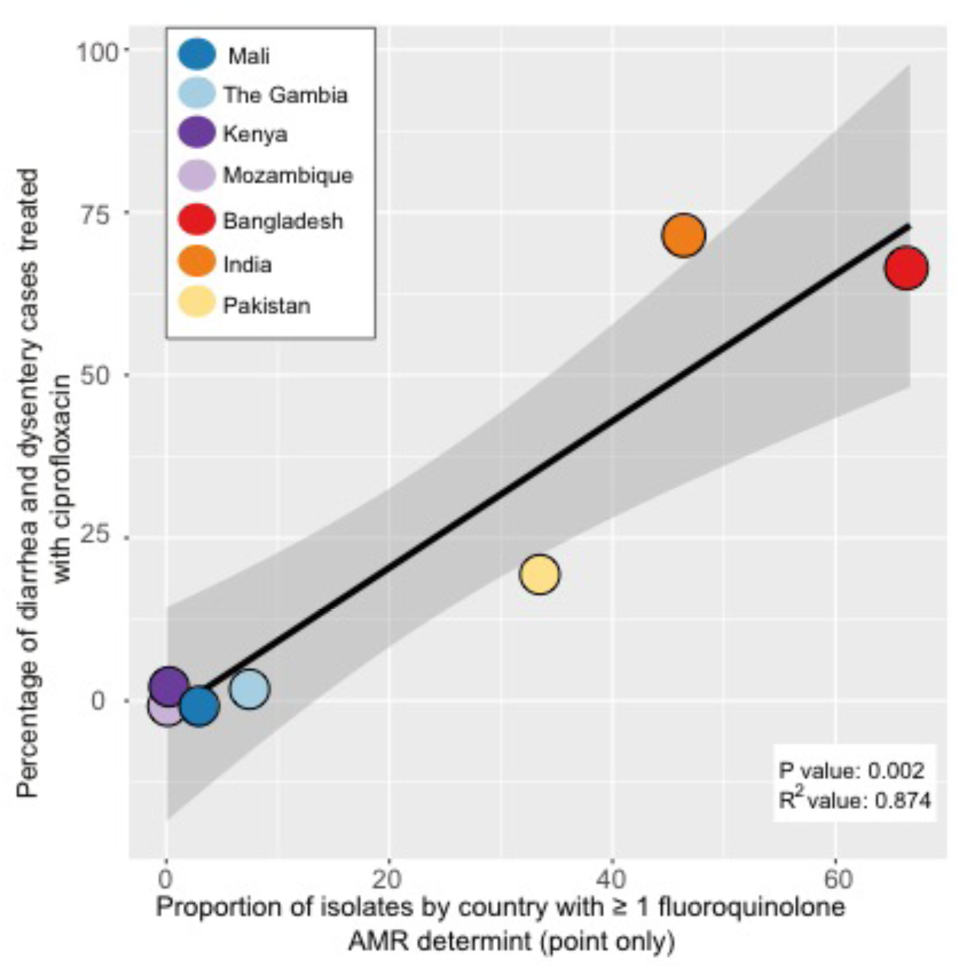
Relationship between use of ciprofloxacin at GEMS study sites and ciprofloxacin resistance in aEPEC. Linear regression (mean ± 95% CI) of proportion of aEPEC with genetic determinants of ciprofloxacin resistance (from point mutations only) vs the proportion of watery diarrhea cases and dysentery case treated with ciprofloxacin at GEMS study sites.

## Discussion

This is the first study to report the prevalence of AMR in community-associated *E. coli* isolates at the GEMS sites. While limited AMR data for *E. coli* are available from the seven countries where the GEMS sites were located, there are no national surveillance data from any of these countries. Furthermore, AMR data on *E. coli* in these countries predominately pertain to *E. coli* causing extra-intestinal infections and to a limited number of drugs, mainly third-generation cephalosporins and fluoroquinolones^21,23-25,31^. Here we present the first comprehensive AMR phenotypic data and molecular characterization for multiple antimicrobials, detecting several common mobile elements mediating the AMR including small plasmids, class I integrons with different cassettes and transposons including *Tn6029, Tn10* and *Tn1721* (Fig S3). Many of the common mobile elements we detected have been reported widely in *E. coli* and other Enterobacteriaceae from human intestinal and extra-intestinal and animal samples^32-36^. Thus, while not conducted for the specific purpose of AMR surveillance, these aEPEC are likely represent a holistic overview of *E. coli,* and possibly other Enterobacteriaceae, in circulation at the study sites. This suggestion is supported by the observation that in this study aEPEC from cases and asymptomatic controls exhibited the same AMR profiles.

The majority (65%) of aEPEC analyzed in this study were MDR, with many displaying resistance to three or more older drugs in key groups: beta-lactams (ampicillin), aminoglycosides (streptomycin), trimethoprim-sulfamethoxazole and tetracycline (Fig 1). The rates of resistance to these drugs generally were lower amongst Asian isolates (21-62%), whereas resistance to newer drugs such as ceftriaxone, fluoroquinolones and azithromycin were detected only at the Asian sites (Fig 3A). These patterns were broadly consistent with antimicrobial usage data from the study sites, which showed that ciprofloxacin and azithromycin were commonly used to treat diarrhea at the Asian sites, while trimethoprim-sulfamethoxazole was the mainstay of treatment at the African sites, supplemented occasionally by other older drugs including ampicillin and chloramphenicol (Fig 3B).

The high AMR frequencies for first-line drugs, in particular penicillins, detected in this study are similar to reports of other intestinal, extra-intestinal and environmental isolates of *E. coli* ^22,37^ Of note, the high levels of ciprofloxacin-resistance found in this study at the Asian sites corresponded to high levels reported in clinical cases in India, Bangladesh and Pakistan ^17,18,21,23,24,38^. Further, while ESBL-producing aEPEC were detected in this study in Asia, the frequency of these was lower than those reported from hospitals in both Asia and Africa^18,21,31,38^. These findings suggest the prevalence of ESBLs, and indeed resistance to carbapenems and last-line aminoglycosides (which were not detected in these community aEPEC) are in higher circulation in hospital settings and have not yet become established in the community settings we investigated.

In agreement with our data relating to AMR phenotypes, we found that the genetic determinants of resistance were similar in bacteria isolated from diarrheal cases and asymptomatic controls (Fig 2A, Table S6) and were not associated with clonal background of the strain, but were associated with the geographic region were the bacteria were isolated (Fig 2B-C; Fig S4). These findings are consistent with the hypothesis that there is selection for different AMR-encoding genes in different locations due to the differing patterns of antimicrobial usage (Fig 3B-C). In strong support of this explanation, mutations in the quinolone resistance determining regions of *gyrA* and *parC* contributed to the regional discrimination function (Fig 2B) and were significantly associated with the frequency of ciprofloxacin use across the seven study sites (Fig 4). However, the situation was more complex for horizontally transferred AMR genes associated with resistance to older drugs. While individual *dfr* alleles were distributed differently across sites (Fig 2C) and contributed to orthogonal components of the regional discriminant function (Fig 2B), the overall prevalence of *dfr* genes was relatively high (50-90%) at each site and not significantly associated with use of trimethoprim (Fig 3D). Similarly, genes encoding resistance to ampicillin, streptomycin and tetracycline were also common in sites where these drugs were rarely used. This could be due to (i) a lack of fitness cost associated with these older resistances, resulting in maintenance of the genes in the *E. coli* population after drug usage declines^11,13,39-41^; (ii) co-selection for resistance to old and new drugs whose associated genes are present in the same mobile elements^42,43^; and/or (iii) selection from drug exposure unrelated to the treatment of diarrhea; for example, the use of trimethoprim to treat other types of infection, or in animal or environmental reservoirs amongst which intestinal *E. coli* circulates. We could not distinguish between hypotheses (i) and (ii) based on our data reported here as we could not determine the precise location of AMR genes from the available short-read sequence data, although further experiments such as conjugation or long-read sequencing could resolve this in future. It is notable, however, that the pCERC-like plasmids (which often carried resistance to streptomycin, sulfonamides and trimethoprim), are very common in *E. coli*, possibly because their small size (~6 kbp, of which half encodes AMR genes) imposes a low fitness cost^32,44^. It is also notable that these and many of the other AMR genes we detected are also associated with composite transposons which can integrate into the bacterial chromosome, where they are maintained at lower fitness cost than large AMR plasmids^36,45,46^.

This study has several key strengths. First, the bacterial samples and usage data were collected seven study sites, four in Africa and three in south Asia, using the same protocols enabling reliable comparisons between the sites^29,47^. Second, the bacterial samples include those from children with diarrheal symptoms and asymptomatic controls in the same study populations, enabling the inference that the aEPEC in this study are broadly representative of the *E. coli* in circulation at these sites. Third, the comprehensive characterization of AMR phenotypes to 16 antimicrobials *in vitro* and their molecular mechanisms from whole genome sequence enabled comparison of both approaches, and demonstrated the value of both types of data. In particular, it showed that reliance on AMR genetic determinants alone may be misleading due to occasional systematic errors in predicting AMR phenotype. Finally, we were able to investigate the complex relationship between antimicrobial use and resistance in community-associated fecal *E. coli* in a way that would be unfeasible using more limited techniques or study sites.

Despite these strengths, there were limitations to this study that could be areas for future improvement. First, the antimicrobial usage data collected at the GEMS sites captured only the drugs prescribed at the GEMS clinics for treatment of diarrheal disease, and does not include antimicrobials available from other sources in the study populations that could also select for AMR *E. coli*, including the treatment of respiratory tract infections^38^. Each GEMS site was expected to follow the WHO guidelines for the management of diarrheal disease^1^, but non-prescribed antimicrobials may account for 19-100% of antimicrobial use outside northern Europe and North America^48^. Increased surveillance and monitoring of antimicrobial use for not only diarrheal disease, but other bacterial infections and their use in agriculture, should help to address this limitation in future. Second, a limitation of the genomic analysis was that the short-read data were insufficient to resolve how all of the AMR-associated genes were mobilized in the aEPEC genomes, which is a common issue in high-throughput genomics studies in general. This could be resolved using long-read sequencing^36,49^, which is becoming more affordable to an extent where it may be feasible to resolve AMR fully in hundreds of bacterial genomes^50^.

## Conclusions

Our data fill important information gaps concerning the prevalence of AMR amongst intestinal *E. coli* circulating in children in seven developing countries in Africa and Asia. In particular, the study showed that MDR involving resistance to multiple first-line or ‘older’ drugs (ampicillin, tetracycline, streptomycin and trimethoprim-sulfamethoxazole) was common at all sites, and resistance to the ‘newer’ antimicrobials, such as fluoroquinolones and azithromycin, had emerged only in Asian sites where these drugs were being used for the treatment of diarrhea. Resistance to older drugs was also common at these sites, such that only Asian isolates were resistant to seven or eight categories of antimicrobials, indicating that changing patterns of antimicrobial use leads to an accumulation of resistance determinants rather than their replacement. Importantly, we found that many AMR genes were encoded on mobile elements that circulate widely amongst *E. coli* and other Enterobacteriaceae, suggesting that these findings may be generalizable to other enteric pathogens and members of the gut microbiome.

## Methods

### aEPEC isolates and corresponding whole genome sequences

A total of 185 aEPEC isolates from the GEMS culture collection were included in the analysis^1,6,29^. Their collection, whole genome sequencing and phylogenomic analysis have been described previously^6^. Briefly, the bacteria were isolated in pure culture from fecal samples from children aged 0-5 years with (cases, n = 94) or without (controls, n = 88) diarrhea at study sites located in seven developing countries (The Gambia, Mali, Kenya, Mozambique, Bangladesh, India and Pakistan)^47^. Three isolates were from children whose case/control status was uncertain. Fecal samples were collected at the study sites before antimicrobial treatment, although prior exposure to antimicrobials from other sources cannot be ruled out. Control children were also not receiving any antimicrobial treatment.

Whole genome sequences were generated for all 185 aEPEC isolates via the Illumina HiSeq platform (100 bp paired-end reads) and assembled using Velvet, as described previously^6,51^. Details of the individual isolates, accessions numbers for the corresponding genome sequence reads and assemblies (deposited collectively under BioProject ERP001141), and associated metadata are provided in Table S1.

### Phenotypic characterization of antimicrobial resistance profiles

Antimicrobial susceptibility testing (AST) to 16 antimicrobials was performed using the VITEK2 (bioMérieux) system or an agar-dilution method. A summary of the drugs, testing methods, and the MIC breakpoints used to determine susceptible (S), intermediate (I) or resistant (R) status for each drug is shown in Table S2. The controls used were three *Salmonella enterica* isolates with known resistance profiles. These were E0001 *Salmonella* Typhimurium PT1 (susceptible to all drugs tested), E004 *Salmonella* Heidelberg (resistant to ampicillin, streptomycin, tetracycline, chloramphenicol, sulfathiazole, trimethoprim, kanamycin, spectinomycin and gentamicin) and E2060 *Salmonella* Thompson (resistant to streptomycin_mod_, tetracycline, kanamycin, nalidixic acid and ciprofloxacin).

For VITEK2 assays, pure isolates were streaked on MacConkey agar plates and grown at 37°C overnight. Isolates were then sub-cultured onto horse blood agar (HBA) plates for fresh culture and grown overnight at 37°C. One to three colonies were selected from each HBA plate and suspended in saline to an OD of ~0.5 MacFarlane Units before being subjected to VITEK2 analysis.

Susceptibility to streptomycin, chloramphenicol, azithromycin, and tetracycline were determined using an agar-dilution method. Bacterial suspensions were first prepared as described above. To each of 32 stainless steel wells, 450 μL of nutrient broth containing 0.05% agar were added, followed by 50 μL of a bacterial suspension. Each Mueller Hinton (MH) agar antimicrobial-containing plate for susceptibility testing and two control MH and MacConkey agar plates were inoculated using a 32-pin replicator. Each pin delivers 2 μL to the plate such that the final number of CFU in each sample was ~10^4^. Plates were incubated overnight at 37°C and inspected the following day. Growth on an antimicrobial-containing MH plate was recorded as phenotypically resistant to the drug, whilst no growth was recorded as susceptible.

The European Committee on Antimicrobial Susceptibility Testing (EUCAST) MIC breakpoints were used where available^52^. Differences exist between the EUCAST and Clinical and Laboratory Standards Institute (CLSI) guidelines in terms of MIC breakpoints and the drugs to be tested. Specifically, tetracycline does not have defined MIC breakpoints under the EUCAST scheme. Hence the CLSI MIC breakpoint was used. Streptomycin and azithromycin do not have established MIC breakpoints under either scheme^53,54^. Previous research proposed a breakpoint of 16 μg/mL for streptomycin in *E. coli*^54^. Little information is available on the MIC distribution of azithromycin for *E. coli*. A breakpoint of 16 μg/mL has been proposed for *Salmonella. enterica* based on a study in which the majority of isolates displayed MIC values of 4 to 8 μg/ml^53^. For the present study, the breakpoint for each of these drugs was set at the conservative MIC of 16 μg/mL. The antimicrobial susceptibility data for each isolate are shown in Table S3.

### Detection of antimicrobial resistance genes

An SRST2-formatted version of the ARG-ANNOT AMR gene database^55^ was downloaded from https://github.com/katholt/srst2. All sequence read sets were screened against the database using SRST2 to detect the presence of known acquired resistance genes in each genome^56^. The beta-lactamase genes, *ampC1*, *ampC2*, *ampH*, were excluded from analysis, as in *E. coli* they are core genes that normally do not confer antibiotics resistance. The results were transformed into a binary table in R to indicate presence/absence of acquired resistance gene alleles (Table S4).

### Detection of SNPS conferring resistance to fluoroquinolones and nitrofurantoin

Chromosomal mutations known to be associated with resistance to fluoroquinolones in *E. coli* were extracted from the genome-wide SNP calls obtained previously based on mapping the reads to the *E. coli* strain 12009 O103:H2 reference genome^6^. These included specific mutations in the QRDR of *gyrA*, *gyrB*, *parC* and*parE*^57^, and non-synonymous substitutions in *nfsA* (residues 11-15) which confer resistance to nitrofurantoin^58,59^.

### Statistical analysis of AMR phenotype prediction from genotype

The ability of genotypes to predict drug susceptibility phenotypes was assessed by comparing summarized AMR phenotypes (S, I and R) with the presence of known AMR-associated genes and mutations. Antimicrobial susceptibility prediction errors are generally characterized as either very major (calling a resistant isolate susceptible) or major (calling a susceptible isolate resistant)^60,61^. The current standard for acceptable very major error and major error rates are <1.5% and <3%, respectively^60^. Here, very major errors were said to have occurred when an isolate was phenotypically resistant, but no known resistance genes or mutations were detected, while major errors were made when an isolate carried known resistance determinant(s) but was phenotypically susceptible. Statistical analysis to determine sensitivity, specificity, positive predictive value (PPV) and negative predictive value (NPV) were calculated in the epiR package for R, using the *epi.stats* function with 0.95 confidence intervals^62^ for each antimicrobial tested.

### Genomic reconstruction of demographic groups by discriminant analysis of principal components

The binary matrix of AMR genetic determinants was used as the input for discriminant analysis of principal components (DAPC) implemented in the adegenet package in R^30^. The first fifteen principal components (PCs), which together explained >95% of the variance in AMR gene content, were retained for discriminant analysis (DA) to explore the ability of PCs to discriminate between groups of strains defined by geographic region of origin (West Africa: The Gambia and Mali; East Africa: Kenya and Mozambique, and Asia: Bangladesh, India and Pakistan) or clonal group membership. The two PCs contributing the most to DA were plotted, and labeled with the genetic determinants whose variation contributed the most to those components. The posterior group membership probabilities for each discriminant function were also plotted.

### Construction of co-occurrence network

A pairwise co-occurrence matrix of acquired AMR genes was constructed by transforming the binary AMR gene content matrix (Table S3) in R. The co-occurrence relationships were visualized between all pairs of genes using pheatmap (Fig S1). Networks of co-occurring genes, in which nodes represent genes and edges represent a frequency of co-occurrence exceeding a given threshold (set to ≥20, ≥33, ≥46, ≥100 genomes), were visualized in R using the igraph package^63^.

### Plasmid replicon screening

An SRST2-formatted version of the PlasmidFinder database^64^ was downloaded from https://github.com/katholt/srst2 for detection of 80 known plasmid replicon marker sequences. All sequence read sets were screened against the database using SRST2 to detect the presence of these replicons in each genome. The results were transformed into a binary table in R to indicate presence/absence.

### Visualization of AMR and plasmid genotypes against a core gene tree

A subtree representing the relationships between the 185 GEMS aEPEC isolates was extracted from the full core phylogeny we published previously^6^ by pruning all other tips using R packages APE^65^ and GEIGER^66^. The presence of acquired AMR genes, mutations and plasmid replicons was plotted as a heatmap against the phylogeny using the plotTree function for R (https://github.com/katholt/plotTree).

### Investigation of mechanisms of AMR gene mobilization

Common AMR-associated genes that were shown to be co-occurring, specifically *bla_TEM-198_*, *sul1*, *sul2*, *strA*, *strB*, multiple *dfrA* alleles, *tet(A)* and *tet(B)*, were further investigated in the aEPEC genome assemblies to determine if they were carried on the same mobile elements. The aEPEC genome assemblies generated previously using Velvet^51^ were interrogated with BLAST, using as queries the AMR genes and the sequences of the plasmids pCERC1 (accession JN012467) and pCERC2 (accession KX291024), and the transposons Tn*6029* (accession CG150541)^45^, *Tn1721* (accession X61367)^67^ and Tn10 (accession AF223162)^68^. For example, if the pCERC2 backbone and AMR genes were all detected on a single contig in the genome assembly, we inferred that these genes were moving together on a pCERC2-related plasmid. Two representative aEPEC isolates that were identified as harboring a pCERC2-like plasmid backbone with different *dfr* gene insertions (strains 402635 and 400879) were selected as representatives for further analysis. These genomes were re-assembled with Unicycler^69^ and annotated using Prokka^70^, and compared to the reference sequences for pCERC1 and pCERC2 using BLAST. The comparison explored using Artemis Comparison Tool^71^ and plotted with genoplotR^72^.

aEPEC isolates were inferred to be likely carriers of Tn6029 or related transposons if the entire region of *repC* to *strB* was detected by BLAST in a single contig and *bla_TEM-198_* was also detected in the genome. (Note it is not possible for the complete transposon sequence to be assembled from short reads, since *bla_TEM-198_* is separated from the rest of the transposon by repeat copies of IS26 which cause a break in the assembly graph). A representative strain matching this pattern (401596) was re-assembled with Unicycler^69^, and the connectivity of Tn*6029* genes in the resulting assembly graphs were visualized using Bandage^73^.

The distributions of class I integrons with different cassette regions were explored by extracting the DNA sequences spanning from *int1* to *sul1* genes using BLAST and MUMmer^74^. Different *dfrA* alleles were identified within the resulting sequences via BLAST searches of the ARG-ANNOT database^55^. Representatives of each distinct class I integron sequence (defined by the *dfrA* gene carried) were re-assembled with Unicycler^69^ and submitted to the Repository of Antibiotic-resistance Cassettes (RAC) website^75^ for detailed annotation.

### Antimicrobial usage data

Data on the use of antimicrobials at each of the seven GEMS sites were collected as part of the original GEMS protocol^1,29^. These data included details of the number and type of antimicrobials prescribed to all cases presenting with watery diarrhea or dysentery at the study clinics. Two of the recorded drugs were excluded from the current analysis: pivmecillinam, because it was not used at any discernible level, and metronidazole, which is active against obligate anaerobic parasites and bacteria only^76^ and therefore is not relevant to *E. coli*, which is intrinsically resistant to this agent. The frequency of prescription of each drug at each site was visualized as a heatmap in R, using the pheatmap package.

The relationship between frequencies of ciprofloxacin and trimethoprim usage and associated genetic determinants was investigated via linear regression modelling in R. For ciprofloxacin, the genetic determinants were either one or more quinolone resistance-associated point mutations in *gyrA* (point mutations in *parC* only occurred when *gyrA* mutations were also present), or the presence of the plasmid-borne genes *qepA* or *qnrS*. Genetic determinants of trimethoprim/sulfamethoxazole resistance were the combination of at least one *dfrA* gene together with *sul1* or *sul2*. The data were visualized in R using the ggplot2 package^30^.

